# Concestor kinase activation mechanism uncovers the cyclin dependence of CDK family kinases

**DOI:** 10.1101/410902

**Authors:** Zahra Shamsi, Diwakar Shukla

## Abstract

Evolution has altered the free energy landscapes of protein kinases to introduce different regulatory switches and alters their catalytic functions. An understanding of evolutionary pathways behind these changes at atomistic resolution is of great importance for drug design. In this work, we demonstrate how cyclin dependency has emerged in cyclin-dependent kinases (CDKs) by reconstructing their closest experimentally characterized cyclin-independent ancestor. Using available crystal structures of CDK2, regulatory switches are identified and four possible hypotheses describing why CDK2 requires an extra intra-domain regulatory switch compared to the ancestor are formulated. Each hypothesis is tested using all-atom molecular dynamics simulations. Both systems show similar stability in the K33-E51 hydrogen bond and in the alignment of residues in the regulatory-spine, two key protein kinase regulatory elements, while auto-inhibition due to a helical turn in the a-loop is less favorable in the ancestor. The aspartate of the DFG motif does not form a bidentate bond with Mg in CDK2, unlike the ancestor. Using the results of hypothesizes testing, a set of mutations responsible for the changes in CDK2 are identified. Our findings provide a mechanistic rationale for how evolution has added a new regulatory switch to CDK proteins. Moreover, our approach is directly applicable to other proteins.

## Introduction

Protein kinases are proteins involved in a variety of cellular signaling pathways that control cell growth. They coordinate cell cycle by switching between active and inactive states, considered as ON/OFF states. Active kinase phosphorylates target proteins to turn ON downstream pathways for signal transduction. Kinase activation is a highly complicated dynamic process, which involves multiple intra- and intermolecular switches that regulate kinase conformational preferences. For example, phosphorylation of the activation loop is one of the most common intra-domain switches regulating the activity of kinases (*1, 2*). These switches are identified using X-ray crystallography, site-directed mutagenesis and computational approaches (*2–7*). The ability of kinases to transfer phosphate groups also depends on an electrostatic network of molecular switches spread across the two lobes. The interaction between two fully conserved residues, glutamate (E) in the *α*C-helix and lysine (K) in the N-terminal lobe is a key switch controlling phosphotransfer (*8*). An aspartate situated in the C-terminal lobe, referred to as a catalytic base, needs to be accessible to the substrate protein to facilitate the extraction of proton from hydroxyl side chains of phosphor-sites of the substrate (*9*). Four hydrophobic residues in the core kinase, called the regulatory spine (R-spine) (Leu66, Leu55, Phe146, and His125, all residue numbers are based on CDK2’s crystal structure (PDB ID: 1FIN (*10*))), align during activation and coordinate the movement of the two lobes (*11*).

In addition to common intramolecular regulatory switches, which can control kinase activation, a variety of intermolecular switches modulate kinase activity via protein-protein interaction. In cyclin-dependent kinases (CDKs), the association of another protein (cyclin) is required for activation. In different cell phases, cyclin can activate CDKs to specifically stimulate distinct signaling pathways (*12*). Similarly, the Src family kinases are also regulated via intermolecular interactions through the SH2 and SH3 domains (*13*). Considering a domino model for the conformational changes in molecular switches that can lead to kinase activation, a series of molecular switches have to become conformationally active along the pathway connecting the inactive and actives state of the kinase domain. If a single switch remains in the “OFF” state, it prevents the overall activation of the kinase. In the past few decades, substantial studies have been carried out to elucidate protein kinase switches in and how they are triggered. However, the question of how different switches work in tandem to regulate the conformational preferences of kinases remains unanswered except for a few well-studied human kinases. Furthermore, it is not clear how these molecular switches have evolved to regulate conformational switching between active and inactive states of kinases.

Proteins with high sequence and structural similarity can display different behavior. Understanding the relation between sequence, structure and function of proteins using large scale molecular dynamics simulations can shed light on how different proteins have evolved conformational regulation. A common way of studying this question is swapping sequences of extant protein and check the functionality to find their relation, which is called horizontal analysis. Hence, we can investigate the effect of changes in each residue, or groups of residues on the protein function. In practice, horizontal analysis of extant proteins has major problems: as the number of suspected residues in the sequence increases and the number of required experiments increases by several orders of magnitude. Due to the highly complicated nature of proteins, these types of methods also experience high frequency of failure in finding sequence, structure and function relations. For example, Src and Abl are two protein kinases 47% sequence identity and a highly conserved three-dimensional structure (*14*). Despite this, they exhibit very different affinities for the cancer drug, imatinib (*15, 16*). Seeliger *et al.*. tried sequence swapping experiments to identify the key residues responsible for the high affinity in Abl or low affinity in Src (*16*). Despite the high sequence identity between Src and Abl, they performed multiple single residue swapping experiments and still could not identify any distinct set of mutations, which could significantly change the Src’s affinity toward imatinib (*16*). Recently, due to advances in sequencing technologies, whole genome sequences for over 1000 species has become available, which makes it possible to reconstruct the phylogeny of modern proteins. This technique, called vertical analysis, makes it possible to follow the changes in residues along evolutionary paths encoding different functional or conformational preferences. Wilson *et al.* applied vertical analysis to answer the challenging question of imatinib selectivity between Src and Abl (*17*). They reconstructed common ancestors of Abl and Src from predicted sequences and tested the drug affinity for each one of them. They also obtained an X-ray crystal structure for one of the ancestors and identified the mechanism of drug selectivity. Another example of successfully finding the sequence-function relation using vertical analysis is study of substrate specificity in CMGC kinase family by Howard *et al.* (*18*). They reconstructed CMGI, the common ancestor of the CMGC family, and tested peptide specificity differences between the CMGI and extant proteins in the family, mainly comprised of CDK kinases (Fig. S1). They observed no co-expression between CMGI and any cyclin or cyclin-like protein even though CMGI was active, which suggests that CMGI, unlike CDKs, is not cyclin dependent. Their observation demonstrates that evolution introduced more regulatory switches in CDKs to make their function more specific. However, the exact set of residues and structural mechanisms responsible for the emergence of cyclin dependency remains elusive.

In this study, we answer the question how cyclin dependency emerged in CDK family kinases using computational ancestral reconstruction of the CMGC family kinase. We study the activation process in CDK2 and the closest common ancestor (concestor) in the CMGC family (named CMGI), which is experimentally proven to be active without co-expression with any cyclin protein (*17*). Then we compare their activation mechanisms in atomistic details to find their differences and similarities and explain how the cyclin dependency has emerged and influenced their activation paths. For the sake of specificity, and considering a significant number of available crystal structures of CDK2, we focus our study on the differences between the modern kinase, CDK2 and CMGI (Fig. S2).

## Results

We hypothesize that in the absence of cyclin, at least one regulatory switch in CDK2 is “OFF” while constitutively remaining “ON” in CMGI. To find suitable candidate regulatory switches, all available crystal structures of CDK2 were compared and four possible molecular switches identified. Based on these switches, four hypotheses are presented.

Cyclin binds to CDKs by forming an interface on with the *α*C-helix and pushing it inward, as observed in the active crystal structure (PDB ID: 1FIN (*10*)). The most intuitive mechanism responsible for cyclin dependence is the rotation/inward motion of *α*C-helix, which can be characterized by hydrogen bonds between Lys33-Glu51 (K-E) and Glu51-Arg150 (E-R). We found that available crystal structures of CDK2 either have formed E-R and broken K-E bonds or broken E-R and formed K-E bonds (Fig. S3). The K-E hydrogen bond is essential for providing electrostatic network required for the process of phosphotransfer while the E-R hydrogen bond facilitates rotation of the *α*C-helix (*19*) (shown in Fig.1 C). Therefore, our “first hypothesis” is whether CDK2 and its ancestor, CMGI, have different equilibrium probabilities of forming and breaking K-E and E-R bonds.

Crystal structures of CDK2 in the inactive conformation exhibit misaligned regulatory-spine (R-spine) residues (Leu66, Leu55, Phe146, and His125) (Fig.1 a&b), suggesting the relevance another regulatory switch (Fig. S4). Cyclin pushes the *α*C-helix inward and leads to the alignment of R-spine residues (Fig.1c). The “second hypothesis” is whether CDK2 and CMGI, have different equilibrium probabilities of R-spine alignment.

**Fig 1:**
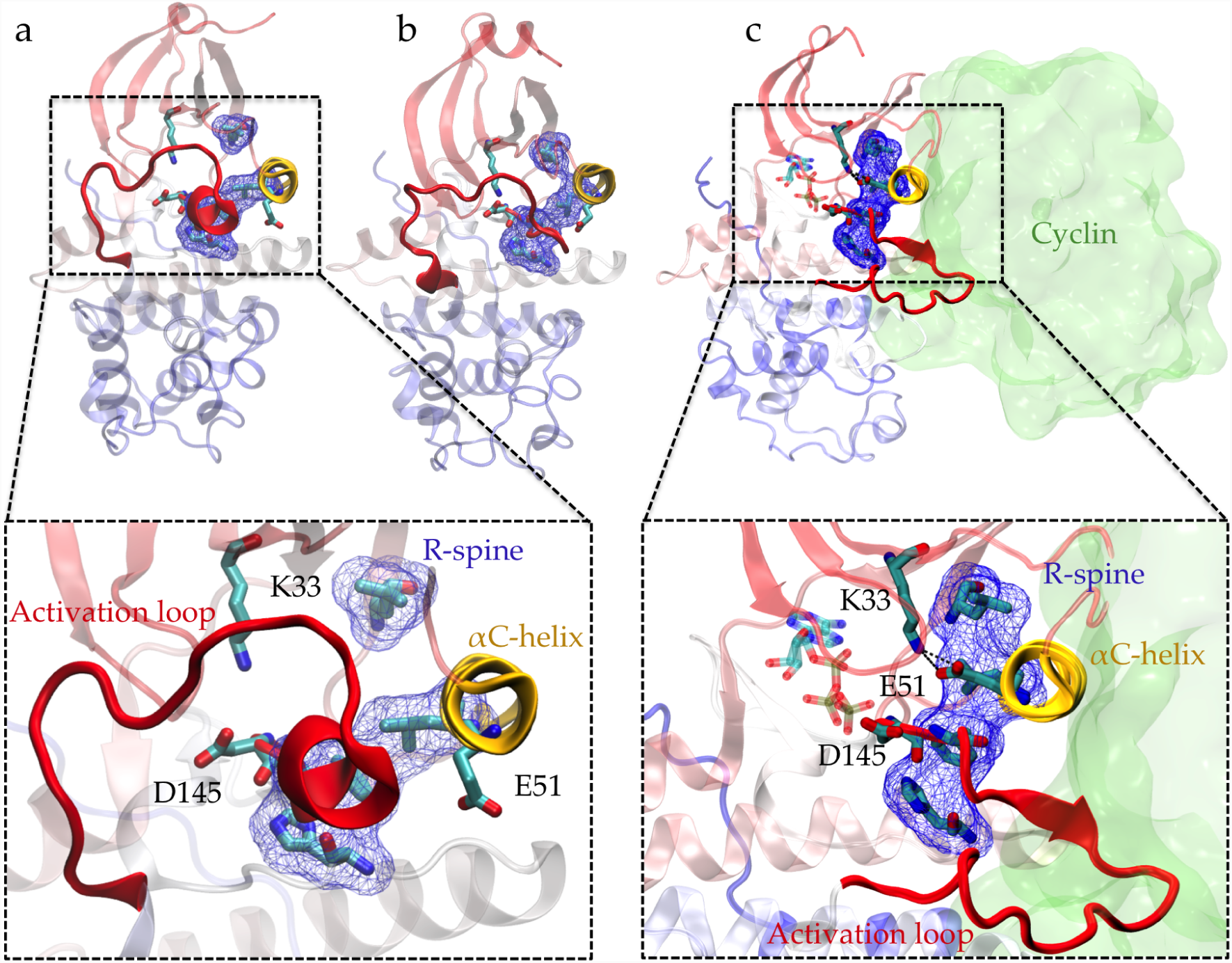
Conformational differences between active and inactive crystal structures of CDK2. Comparison of the (a, b) inactive (PDB IDs: 3PXF (*20*) and 4GCJ (*35*)) and (c) active crystal structures (PDB ID: 1FIN (*10*)) highlights the conformational changes associated with activation process; activation loop (a-loop) in red adopts different folded conformation, *α*C-helix in yellow rotates, electrostatic network formed between K33, E51 and D145 switches and alignment of residues Leu66, Leu55, Phe146, and His125 known as regulatory spine (R-spine) (shown in licorice and blue surface representation) alters. In the active crystal structure (c) cyclin is also shown (with green surface representation) in its bound position next to *α*C-helix.

Formation of a helical region in the beginning of activation loop (a-loop) is another characteristics of inactive CDK structures, which prevents binding of the substrate protein (PDB ID: 3PXR and 3PXF (*20*)). The helical turn pushes the *α*C-helix out, thereby acting as a molecular switch that could alter the cyclin dependence of CDK kinases (Fig.1). This auto-inhibitory mechanism is observed in several kinases such as CDKs, Src and Abl (*21*). Crystal structure analysis shows high degrees of correlation between the presence of a helical turn and existence of the K-E bond (Fig. S5). Possible differences in helical turn formation between CDK2 and CMGI is our “third hypothesis”.

The precise orientation, and positioning of the triad of highly conserved residues, Lys33 (K33), Glu51 (E51) and Asp145 (D145), is crucial for catalysis and phosphotransfer processes in kinases (*10, 22*). Orientation of Asp145 (in the well-known DFG (Asp145, Phe146 and Gly147) motif) (Fig.1) is particularly critical due to its interaction with Mg^2+^ ions, serving as a shuttle for cations to ATP phosphate groups (*10, 22–24*). Even though the exact catalytic role of Asp145 is not well understood, some crystal structures of cAMP-dependent protein kinase (another family of kinase) captured the phosphoryl transfer. These crystal structures show Asp145 forms a bidentate bond with one of the Mg^2+^ ions, which is another regulatory switch in protein kinases (*25–28*) (see Fig. S13). Previous quantum mechanical calculations also show that Asp145 forms a bidentate bond with a Mg^2+^ ion in its active structure (*29*). As there are no Mg^2+^ ions in most of the crystal structures, availability of Asp145 is measured by calculating Asp145-Lys33 distance in crystal structures of CDK2, which suggest existence of two distinct states in the space of K33-E51 (Fig. S6). Cyclin binding/unbinding can alter the orientation, accessibility and hydrogen bonds formed by Asp145 (*12*) in CDK2, while in CMGI they may get aligned without cyclin binding. This can be the difference between CDK2 and CMGI, which is our “fourth hypothesis”.

In this study, we test each hypothesis by investigating the dynamics behavior of the switches in CDK2 and CMGI via large timescale unbiased molecular simulations (see Materials and Methods).

### K-E hydrogen bond is equally stable in CMGI and CDK2

Comparison of free energy landscapes in Fig.2 shows the relative free energies of the formed K-E bond state (active-like state, where the distance between Lys33 and Glu51 is less than 0.5 nm) is comparable between CDK2 and CMGI, suggesting that they have similar stabilities (Fig.2 region D and *δ*), which refutes our first hypothesis (see Fig. S7 for raw distributions).

**Fig 2:**
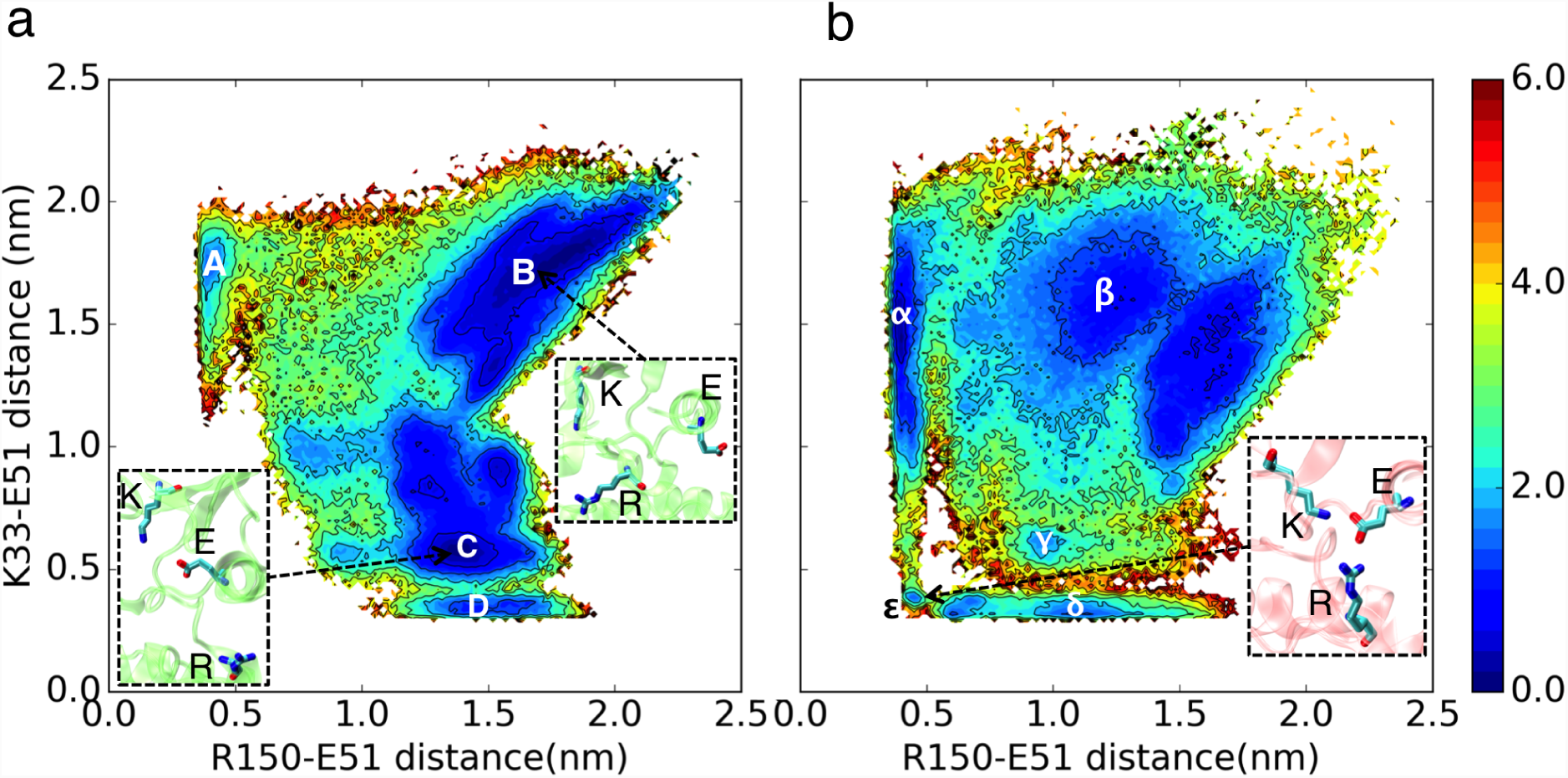
Comparison of free energy landscapes of E-R versus K-E distances between CDK2 (a) and CMGI (b). Colors show the free energy in kcal/mol.

The interaction between Arg150 (residue in the a-loop) and Glu51 (E-R) is a competitor for Lys33-Glu51 (K-E) bond. The presence of an intermediate state with both the K-E and R-E bonds formed (Fig.2 region ∊, with K-E *<* 0.5 nm and E-R *<* 0.5 nm) in CMGI reveals an alternative activation pathway where an intermediate state, with triplet Lys33, Glu51, and Arg150 interacting, facilitate formation of the K-E bond. This facilitation process has been observed in other kinases, including other CDKs, in previous studies (*19, 30*). In the landscape of CDK2, the K-E bond forms only when the E-R bond is broken consistently with the lack of evidence for a triple interaction in CDK2.

### R-spine acts similarly in CMGI and CDK2

The conformational landscape of CDK2 in Fig.3 does not display any considerable barrier for alignment or misalignment of R-spine (there is no high energy region to move along x direction in Fig.3). In both conformational landscapes, R-spine can form or break easily when the K-E bond is broken (when K-E is larger than 0.5 nm, RMSD of R-spine can be either high or low), whereas when K-E bond is formed the R-spine does not break (when when K-E is less than 0.5 nm, RMSD of R-spine with respect to the active structure is always low). The similar dynamic behavior between the R-spine of CMGI and CDK2 disqualifies our second hypothesis (see Fig. S8 for raw distributions).

**Fig 3:**
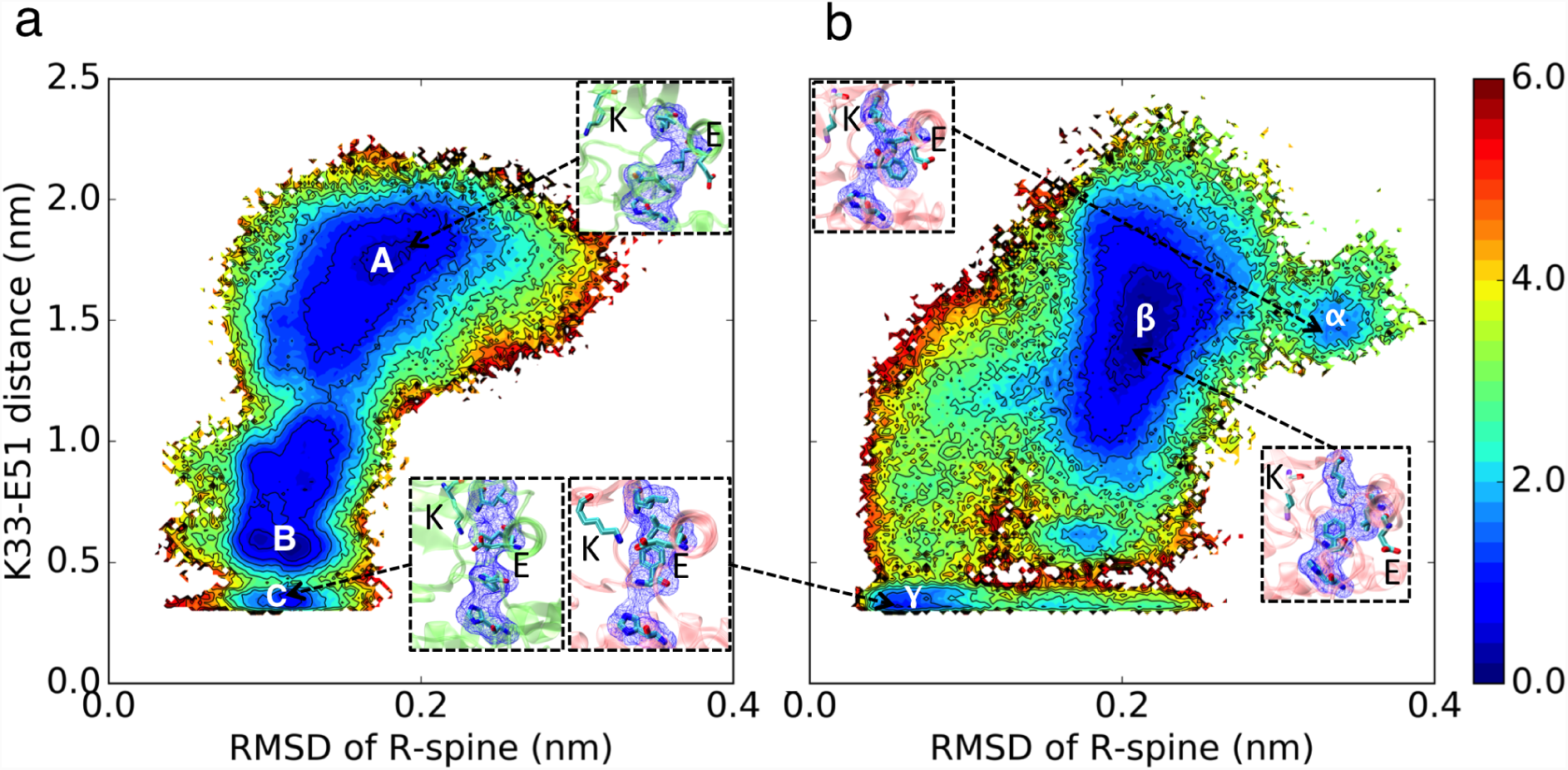
Comparison of free energy landscapes of R-spine RMSD versus K-E distances between CDK2 (a) and CMGI (b). Colors show the free energy in kcal/mol. R-spine RMSDs were calculated with respect to active crystal structure (PDB ID: 1FIN (*10*)) in CDK2 and active structure from simulation in CMGI.

### Auto-inhibition due to the helical turn in the a-loop is less probable in CMGI compared to CDK2

The simulation results projected onto two dimensional conformational landscape of K-E distance versus RMSD of helical turn in the beginning of a-loop with respect to inactive structure (PBD ID: 3XPR (*20*)) reveals a barrier of 6 kcal/mol for unfolding of helical turn in CDK2, whereas the barrier is not observed due to an stable intermediate state in CMGI (in Fig.4, region *β* is the intermediate state in CMGI, which its low free energy facilitates the transition from inactive state *α* to active-like state *δ*, while respectively region **B** in the CDK2’s landscape is less stable and makes a barrier of 6 kcal/mol for the transition from inactive state **A** to active-like state **D**) (see Fig. S9 for raw distributions). The helical secondary structure moves from beginning of the a-loop toward its end in the *β* intermediate state in CMGI (Fig.4 b), which allows the a-loop to fully unfold with a lower barrier. These free energy landscapes support our third hypothesis that differences in the stability of the a-loop helical turn between CDK2 and CMGI are one of the factors distinguishing their cyclin dependence. However, the origin of the different in helical turn stability remains unclear.

**Fig 4:**
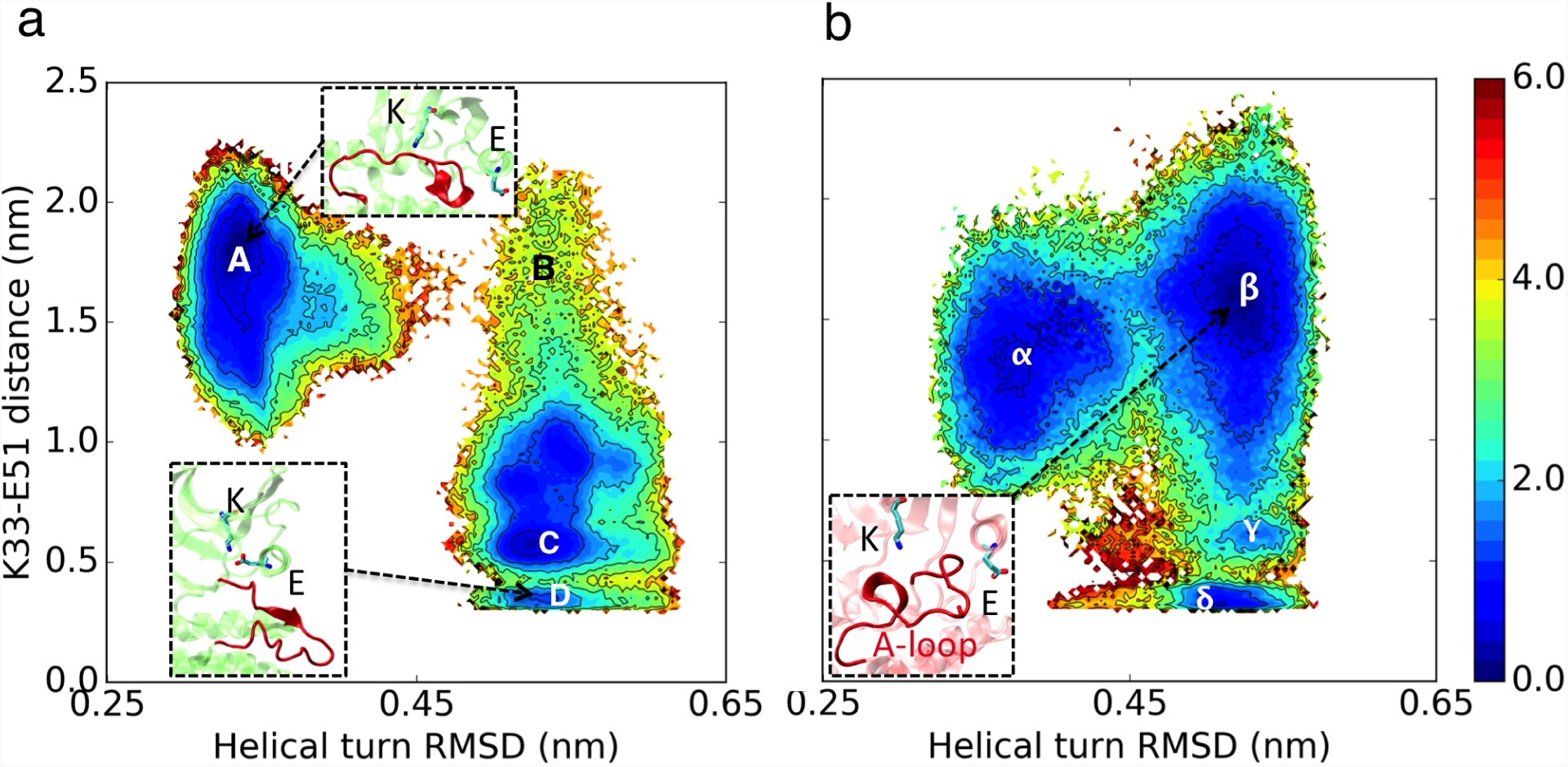
Comparison of free energy landscapes of helical turn RMSD versus K-E distances between CDK2 (a) and CMGI(b). Colors show the free energy in kcal/mol. Helical turn RMSDs are calculated with respect to inactive crystal structure (PDB ID: 3PXR (*20*))

Crystal structural analysis of the CDKs reveals there is a salt bridge between His161 in the a-loop and Glu12 in the p-loop in all inactive CDK2 crystal structures with helical turn (Fig. S10). This ionic interaction stabilizes the upward (when the a-loop is closer to the N-terminal lobe, shown in Fig. S10. 1) conformation of the a-loop, which provides enough space for formation of the helical turns. His161 in CDK2 is substituted with Glu161 in the ancestor. Repulsion between Glu12 and Glu161 destabilizes the upward conformation of the a-loop and consequently prevents formation of the helical turn (Fig. S11 & S12). The free energy landscape of K33-E51 versus E12-H161 shows that the 3 kcal/mol barrier for unfolding of the helical turn in CDK2 is due to E12-H161 bond breaking. Another reason for the difference in the helical turn stability can be due to their differences in the helical turn amino acid sequences.

### The aspartate in DFG motif does not form a bidentate bond with Mg^2+^ in CDK2

In our simulations, Asp145 can interact with a Mg^2+^ ion with two different bond types: it can form a bidentate bond between its two carboxyl oxygens and Mg^2+^ or a single bond between one of its carboxyl oxygens and a Mg^2+^ ion (Fig.6 and 5) (see Fig. S14&S15 for raw distributions). Based on the simulation results, CDK2 without cyclin bound has a very low probability of having K-E bond and bidentate D145-Mg^2+^ bonds at the same time, which is the property of observed active structures (Fig.6 region **D**). In contrast, the CMGI the low free energy state *δ* (Fig.6) has both bidentate D145-Mg^2+^ and K-E bonds. In both systems, D145 can form both types of interactions with Mg^2+^ while the K-E bond is broken.

**Fig 5:**
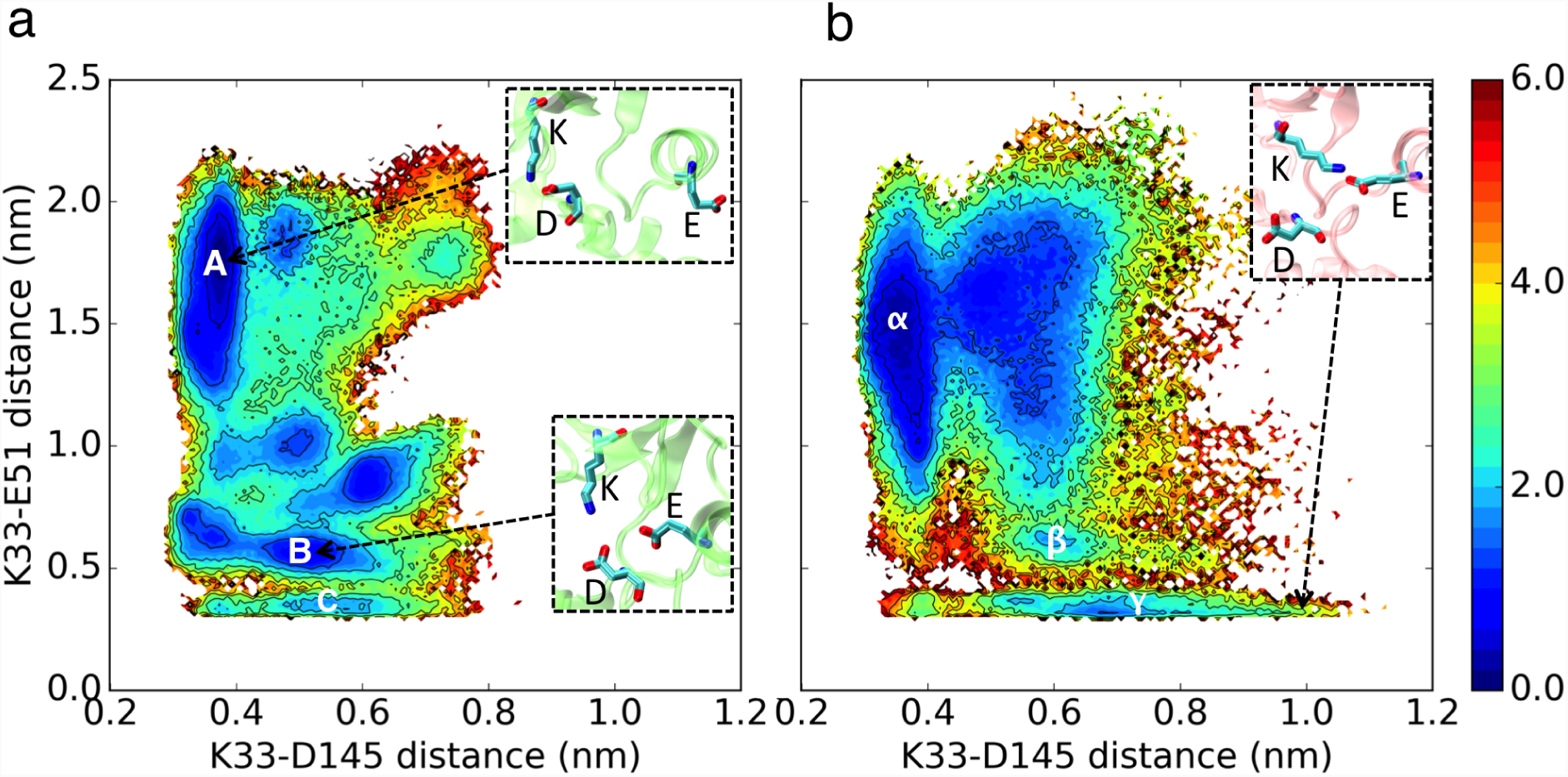
Comparison of free energy landscape of K-D versus K-E distances between CDK2 (a) and CMGI(b). Colors show the free energy in kcal/mol.

**Fig 6:**
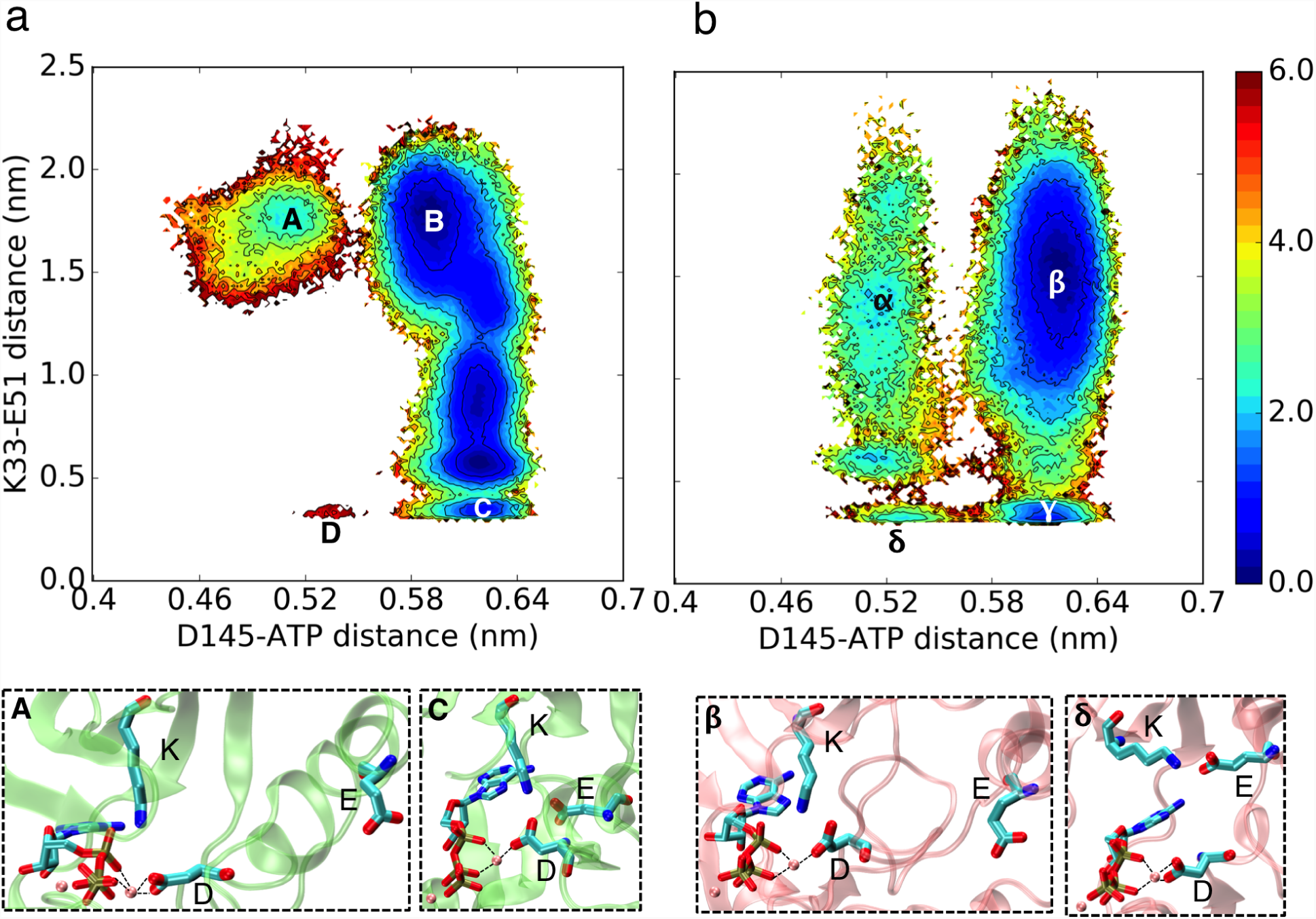
Comparison of free energy landscape of ATP-D versus K-E distances between CDK2 (a) and CMGI(b). Colors show the free energy in kcal/mol.

In order to determine, which type of Asp145-Mg^2+^ bond active, cyclin-bound CDK2 forms, an additional 2 *μ*s simulations of CDK2 bound to cyclin with ATP and two Mg^2+^ ions were performed These simulations show a significantly different stability profile for Asp145-ATP distance between cyclin-bound and cyclin-free CDK2. Unlike the CDK2 monomer, the CDK2-cyclin dimer demonstrates no barrier for switching between the two bond types, which is similar to CMGI. Cyclin binding changes the electrostatic network of the residues in a way that makes the APS145-Mg^2+^ bidentate bond more stable (see Fig. S16).

### CMGI activates through a different path compared to CDK2

Transition Path theory (TPT) analysis indicates that CDK2 goes through more states to activate compared to CMGI. As the probability of bidentate bond in Asp145-Mg^2+^ along with other regulatory switches is negligible in CDK2, in the TPT studies, we disregard the Asp145-Mg^2+^ bond and focus our study on other regulatory switches. TPT was used to analyze the activation path with highest flux from inactive to active-like states (Fig. S27 and Fig. S28). In the activation pathway of CDK2, strong formation of R-spine and helical turn prevents *α*C-helix from rotating and the K-E bond from forming. As the helical turn unfolds, lysine finds space to push the *α*C-helix inward and rotate it to form the K-E bond. The processes of the R-spine breaking and re-forming, the K-E bond forming and unfolding of the helical turn, happen in a single MSM state transition, while the K-D bond breaking occurs in another MSM state transition. Unlike CDK2, in CMGI the helical turn unfolds first, and then in the next MSM state transition R-spine forms simultaneously as K-E bond. K-D bond breakage also has a lower free energy barrier in the CMGI.

## Discussion

Our simulations confirm the experimental observation that the CDK ancestor, CMGI, is active in the absence of cyclin, unlike the cyclin-dependent CDK2. All CMGI’s regulatory switches can be in the “ON” mode at the same time independent of any intra-molecular interactions, whereas two of the regulatory switches of CDK2, K-E bond Asp145-Mg^2+^ bond can not be in the “ON” mode simultaneously. Folding of the helical turn in the beginning of the a-loop, blocks the substrate binding path in CDK2 without cyclin. The critically important conserved Asp145 residue also displayed different behavior in the CDK2 monomer compared to the CDK2-cyclin dimer and CMGI. The preferred orientation of Asp145 in the monomer CDK2 tightly bound to Lys33 not only makes the residue less accessible to the substrate but also prevents the formation of a bidentate bond with Mg^2+^ ion, reducing kinase activity.

Homogeneous active versus heterogeneous inactive state is observed in kinases. While a defined active conformation is needed to guarantee a catalytically competent active site and specific interaction with downstream partners, deactivation of a protein kinase can be accomplished by a shift to any conformation other than the active structure. The intrinsically entropic nature of the inactive state may not be a limitation in the efficiency of the conformational transition, but rather provides an advantage. Modern proteins have evolved mechanisms for enhanced regulation. Evolution has significantly changed features of free energy landscapes and created more complex landscapes by introducing multiple new minima (Fig. 5). It also demonstrates that CDK2 spends less time in the regions, which are functionally unimportant.

A visual inspection of the kinetic plots suggests that thermal fluctuations toggle all the molecular switches via a concerted mechanism, where molecular switches are triggered cooperatively. However, looking at the conformational landscape of molecular switches reveals a different, more sequential view of CDK activation, where some molecular switches are turned ON/OFF before other switches change their conformation. This observation is directly related to a prominent debate about conformational change mechanisms in general, comparing a sequential domino brick effect or Monod-Wyman-Changeux type of concerted action allostery (*31–34*) (Fig. S17 and S18 show the one-dimensional probability density map for each metric) (Fig S19 to S26 show two-dimensional probability densities and free energies for combination of each two of the metrics).

Our study raises several interesting questions about the evolution of protein structure and function: 1) How network of protein conformations evolves to acquire a specific function or integrate an external signal in the form of binding partners such as ligands and other proteins? The process could be elucidated by investigating ancestral proteins along the evolutionary trajectory. However, the large computational time requirements would make such an investigation intractable, which calls for development of more efficient computational approaches to enable computational ancestral protein reconstruction. 2) How the conformational network properties change during evolution? These properties include network connectivity i.e number of connections per state and robustness i.e. how many states and edges (connections between states) could be removed without altering the overall function. 3) Are modern signaling proteins more efficient than ancestral proteins in terms of energy dissipated during functional dynamics? 4) Finally, the question of how functional free energy and folding free energy landscapes are designed during evolution to enable protein conformational change while keeping it in the folded state?

## Materials and Methods

### Generation of initial structures using Modeller and accelerated molecular dynamics simulation

One active (PDB ID: 1FIN (*10*)) and three different inactive (PDB IDs: 3PXR (*20*), 4GCJ (*35*) and 3PXF (*20*)) X-ray crystal structures of human CDK2 kinase were used as starting structures for CDK2. As kinases have well conserved three dimensional structures, we used the CDK2 crystal structures as templates for modeling CMGI. Using the Modeller (*36*) with CDK2 crystal structures (PDB IDs: 1FIN (*10*), 3PXR (*20*), 4GCJ (*35*) and 3PXF (*20*)) as templates, four homology models were built for CMGI as starting structure.

In each system, molecules except CDK2 (or CMGI) were removed. Phosphate on Thr160 and an ATP molecule with two magnesium ions bound, taken from previous simulations, was inserted into the binding pocket. The starting structures were solvated in water boxes, with dimensions of approximately 85 Å X 70 Å X 60 Å with TIP3P model molecules (*37*). Sodium and chloride ions were added to neutralize the charge of all systems and bring salt concentration to approximately 150 mM. All systems were subjected to 10,000 steps of energy minimization and were equilibrated for 2-4 ns in an NPT ensemble at 300K and 1 atm. Simulations were performed using a 2 fs time step, periodic boundary conditions, and constraints of hydrogen-containing bonds using the SHAKE algorithm (*38, 39*).

Equilibrated structures simulated for one set (5 *μ*sec for each system) using accelerated molecular dynamics (aMD) as a preliminary simulation to obtain starting structures for the production unbiased MD runs (*40, 41*) (see Supporting information for aMD parameters). Starting structures for unbiased production run were chosen randomly from landscapes covered by aMD.

All simulations run on CUDA version of AMBER 14 (*42*), using AMBER14 forcefield, ff14SB for proteins (*43*) and General AMBER Force Field (GAFF) (*44*) for ATP on the Blue Waters supercomputer. Total aggregated unbiased MD of 76 *μ*sec for CDK2 and 42 *μ*sec for CMGI were performed.

### Markov state models

Markov state models (MSMs) are kinetic models to model randomly changing systems like protein dynamics (*45*). An MSM represents protein dynamics as a Markov chain on discretized conformational space achieved by clustering of protein conformations in MD trajectories. Transitions between discretized states in MD trajectories are counted and a transition probability matrix is estimated using the maximum likelihood method. If vector *p*(*t*_0_) denotes the probability of being in any of the states at time *t*_0_, the probabilities at time *t*_0_ + *kτ* are given by:

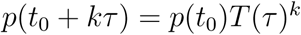

where *T* (*τ*) is the transition probability matrix parameterized by a lag time, *τ*. MSMs accurately approximate protein dynamical processes with timescales relevant to biomolecular function, far longer than any individual trajectory used in MSM construction (*46–48*).

Adaptive sampling is a computational technique to enhance the simulation of biomolecular functions and folding. Adaptive sampling involves iteratively running short simulations, clustering on a relevant metric, and seeding new simulations from clusters based on some criterion. Adaptive sampling has been shown to sample configurational space more efficiently than the simulated tempering method for simulation of an RNA hairpin (*49*). MSMs importantly estimates the equilibrium populations of states from trajectories sampled from non-equilibrium distributions and generate unbiased transition probabilities, allowing for accurate characterization of both kinetics and thermodynamics.

In order to build MSMs, the system’s dynamics should discretize into a relevant metric. We calculated root mean square fluctuations (RMSF) of all residues to identify residues with higher fluctuations, which shows that they participate more in the kinase dynamics (see Fig. S29 and Fig. S30) (residues 31 to 83 and 145 to 177 in CDK2 and 31 to 100 and 145 to 180 in CMGI). Based on the literature, we knew that residues in the C-lobe are not participating in the activation process, so did not include them even with high RMSF values. Dihedral angles (*ϕ* and *ψ*) of these residues were considered as raw features. Time-structure independent component analysis (tICA) was used to reduce the dimension of the high dimensional dihedral angle metrics space by projecting onto the slowest subspace (*50, 51*). To build optimal MSMs, we varied the numbers of clusters along with numbers of tICA components to project onto in order to build our MSMs. The generalized matrix Rayleigh quotient (GMRQ) (*52*) score and percentage of the used data were calculated for each MSM and parameters, which give the higher GMRQ score with the higher data usage were picked as the best sets (see Fig. S31 and Fig. S32) (1000 clusters with 10 tICA components for CDK2 and 300 clusters with 6 tICA components for CMGI were picked as the best set). To find the best lag time, series of MSMs with different lag times were built and the implied timescales were calculated to find a region where the spectrum of implied timescales were relatively insensitive to lag time. A lag time of 14 ns was suitable for both systems (Fig. S33 and Fig. S34).

All MSM analysis in this study was conducted using the MSMBuilder3.8.0 Python package (*53*).

### Transition path theory

Transition path theory (TPT) is a rare event sampling method allows for the determination of the likelihood of transition along with pathways in the Markov random field between the two states. We used the MSMBuilder implementation of TPT in order to identify top pathways from the net flux matrix. For a detailed overview of TPT, we refer the reader to a recent review by Vanden-Eijnden et al. (*54*).

## Acknowledgements

### Competing Interests

The authors declare that they have no competing financial interests.

### Data and materials availability

Additional data and materials are available online.

